# Short T1 Fraction as a Marker of Myelin Content: Evidence from Postmortem MRI and Histology

**DOI:** 10.64898/2026.05.31.728778

**Authors:** Chenyang Li, Dominique Leitner, Zifei Liang, Veronika Yakovishina, Merna Ibrahim, Arline Faustin, Thomas Wisniewski, Youssef Zaim Wadghiri, Yulin Ge, Jiangyang Zhang

## Abstract

**Purpose:** To investigate the short-T1 fraction as a potential biomarker of white matter myelin integrity, using myelin histology as ground truth.

**Methods:** Multi-inversion-time MRI data were acquired from four postmortem brain hemisphere specimens from donors with Alzheimer’s disease on a clinical 3T scanner, and from five dissected tissue blocks containing white matter hyperintensities (WMHs) on a preclinical 3T system. Short-T1 fraction maps were generated using inverse Laplace transform to isolate short-T1 components and were compared with T2-based myelin water imaging metrics through joint T1-T2 correlation analysis. Short-T1 fraction maps from the tissue blocks were further compared with myelin-stained histology. In addition, *in vivo* short-T1 fraction data were acquired from two elderly volunteers with WMHs to demonstrate translational feasibility.

**Results:** Postmortem MRI revealed reduced short-T1 fractions in WMHs. T1–T2 correlation analysis showed that the short-T1 fraction was closely associated with the short-T2 (myelin water) component. Strong correlations were observed between short-T1 fractions and optical densities from both Luxol Fast Blue- and myelin basic protein-stained histological sections, supporting the link between the short-T1 signal and myelin content. Consistent findings were also observed *in vivo, where* significant reductions in short-T1 fractions were detected within WMHs.

**Conclusion:** The short-T1 fraction correlated with myelin content in postmortem brain white matter, supporting its potential as a clinically translatable biomarker of myelin integrity.

## 1. Introduction

White matter integrity is a key determinant of cognitive function and brain health across the human lifespan. Progressive white matter degeneration is commonly observed in aging and Alzheimer’s disease (AD) and AD-related disorders(1,2). For example, white matter hyperintensities (WMHs) are closely associated with vascular risk factors and chronic inflammation in aging and AD(3–6). Myelin pathology is a common component of WMH pathology(5,7), and noninvasive assessment of myelin integrity in the brain is important for monitoring the axonal degeneration and progression of aging and AD.

Several MRI techniques have been developed for characterizing myelin integrity in the brain. Among them, relaxometry-based MRI methods have been widely used to probe myelin’s unique microstructural and biochemical properties. Myelin water imaging (MWI), which quantifies the short-T2 signal component(8,9), presumably from water located between myelin bilayers, has shown strong correlations with histological measures of myelin(10). Similar T2*-based approaches have also been developed to map myelin water fractions (11–14). A detailed overview of MWI techniques can be found in prior reviews(15–17).

More recently, T1-based approaches have gained attention as a new strategy for quantifying myelin water fraction. This is based on the assumption that myelin water has a shorter T1 than other cellular water compartments. For instance, Labadie et al.(18) applied spectral analysis of multi-inversion-recovery T1 data to separate short- and long-T1 components, with the short-T1 fraction presumed to reflect myelin water due to its lipid-associated environment. Several T1-based myelin water imaging (MWI) techniques have been developed for potentially faster whole-brain myelin mapping compared with conventional T2-based MWI methods. Because T1-based measurements primarily rely on the selection of longitudinal magnetization rather than transverse signal evolution, they minimize the stimulated echo contamination(19) and mitigate the tradeoff between short echo spacing and high spatial resolution that commonly limits T2-based MWI techniques. For example, double inversion recovery (DIR) and triple inversion recovery approaches have been proposed to suppress water signals from multiple tissue types while preserving the short-T1 myelin water signal (20–23). Oh et al.(24,25) further developed the direct Visualization of Short Transverse Relaxation Time Component (ViSTa) technique, which uses a DIR scheme with a shorter repetition time to highlight short-T1 signals while suppressing long-T1 signals. Ma et al.(26–29) proposed short TI adiabatic inversion recovery (STAIR), which uses a single steady state inversion recovery acquisition to attenuate long-T1 signals while preserving short-T1 signals. Moreover, the multicomponent driven equilibrium single pulse observation of T1 and T2 (mcDESPOT) approach was proposed for myelin water imaging based on the same assumption (30,31). Although these short-T1 imaging techniques are promising, the exact relationship between the short-T1 fraction and underlying tissue myelin content is still not fully understood, due to limited studies with direct comparisons with histopathology. Establishing this relationship will be essential for the further development of these techniques for diagnosis of myelin injury in WMHs and other myelin-related pathologies.

In this study, we investigated whether the short-T1 fraction can serve as a robust marker of myelin-related changes in WMHs. We acquired multi-inversion time MRI data from postmortem brain specimens from patients diagnosed with AD with WMHs followed by histopathological evaluation. We then evaluated its relationship with myelin content from histology. As a proof of concept, we further demonstrated *in vivo* mapping of short-T1 fractions in subjects with WMHs.

## 2. Method and Materials

### 2.1 Brain specimen sample preparation

Four human brain specimens with postmortem diagnoses of AD were obtained from the Alzheimer’s Disease Research Center (ADRC) at NYU Grossman School of Medicine (Table 1). Neuropathological assessments were performed by ADRC according to the National Institute on Aging–Alzheimer’s Association (NIA-AA) criteria(32). Among the four donor cases, three demonstrated high Alzheimer’s disease neuropathologic change (A3B3C3), while one showed intermediate AD neuropathologic change (A2B2C2). Informed consent was obtained for all cases for their in vivo clinical research studies, and all procedures were approved by the NYU Grossman School of Medicine Institutional Review Board. Specimen details, including postmortem interval (PMI), fixation duration, and pathological diagnosis, are summarized in **Table 1**. All experiments were performed in accordance with NIH guidelines and regulations for postmortem analysis.

**Table 1.**
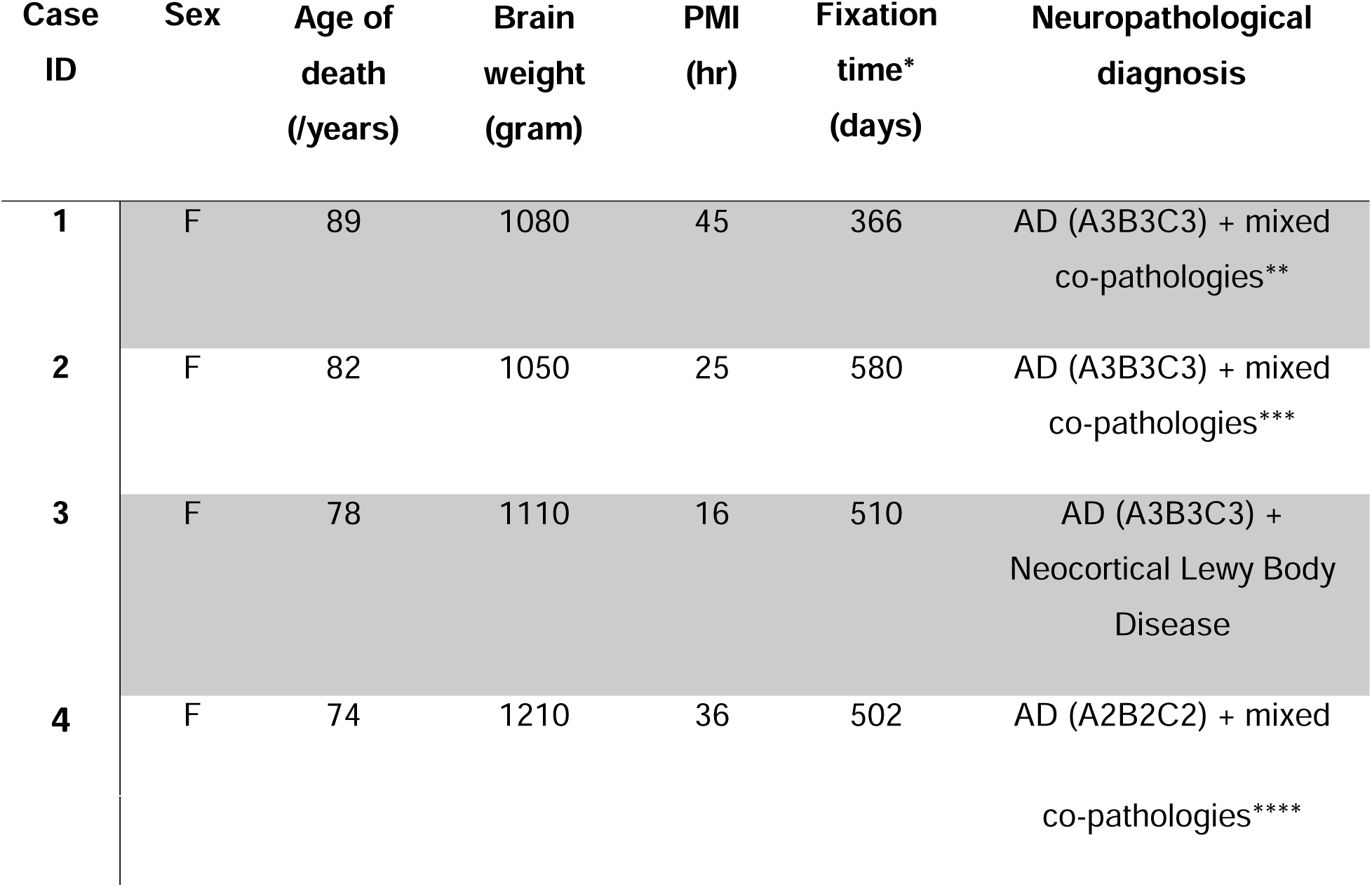
Demographic and pathological information for brain specimens. *Fixation time was calculated from the date of autopsy to the days of MRI scanning. ** Co-pathologies for Case 1 include remote amygdala infarct, hippocampal sclerosis, Binswanger disease, arteriolosclerosis, moderate CAA, amygdala-predominant LBD, LATE-NC stage 2, and ARTAG. *** Co-pathologies for Case 2 include mild cortical atrophy, mild atherosclerosis, mild WM rarefaction. **** Co-pathologies for Case 4 include mild atherosclerosis, mild WM rarefaction, limbic LBD, and widespread TDP-43 pathology.

Prior to MRI, the hemisphere specimens were stored at 4C^0^ in phosphate-buffered saline (PBS) for two weeks with fresh PBS every 3-4 days to washout excess formalin and reverse the changes in tissue relaxation times due to formalin fixation (33). One day before MRI, each hemisphere was transferred to proton-free Fomblin (perfluoropolyether) and fully submerged in a sealed plastic container to minimize susceptibility artifacts. The specimens were subsequently placed under vacuum to remove trapped air bubbles. After MRI of the hemispheres on a clinical 3 Tesla (T) Prisma scanner, specimens were rinsed in PBS and returned to the neuropathology core for further dissection. Locations of WMH lesions were identified based on MRI findings, and corresponding tissue blocks (approximately 40 mm x 28 mm x 5 mm) were dissected and placed in pathology cassettes (**Supplementary Figure S1**). The cassettes were immersed in PBS within an 80-mL syringe for subsequent MRI examination on a Bruker 3T scanner, followed by histopathological analysis.

### 2.2 Postmortem MRI protocols

#### 2.2.1 MRI of postmortem brain hemispheres on a clinical 3T system

Postmortem brain hemispheres were imaged on a clinical 3.0 T Prisma (Siemens Magnetom) using a 64-channel head coil. The MRI protocol included the following sequences:

(**a**) **3D T1-weighted Magnetization Prepared Rapid Gradient Echo (MPRAGE)**: Images were acquired with flip angle = 8°; inversion time (TI)/echo time (TE)/repetition time (TR) = 500/2.14/2100 ms; matrix size = 256 × 256 × 160; isotropic resolution = 0.8 mm. The inversion time (500 ms) was empirically chosen to yield gray–white contrast comparable to *in vivo* MPRAGE, accounting for fixation-related relaxation changes in postmortem tissue.
(**b**) **3D T2-FLAIR**: Images were acquired with the following parameters: TI/TE/TR = 1100/394/4000 ms; matrix size = 256 × 256 × 160; isotropic resolution = 0.8 mm. The TI was selected to suppress signal from PBS.
(**c**) **3D T2-weighted SPACE**: Images were acquired with TE/TR = 104/3000 ms; matrix size = 256 × 256 × 160; isotropic resolution = 0.8 mm, and were used to complement FLAIR images for delineation of WMH lesions.
(**d**) **Short-T1 MRI**: We used an inversion-recovery–prepared turbo spin-echo (IR-TSE) sequence with 19 TIs (0, 25, 50, 100, 150, 200, 250, 300, 350, 400, 500, 600, 700, 800, 900, 1000, 1200, 1600, 3500 ms), effective TE/TR = 8.5/8000 ms, turbo factor = 4, matrix size = 256 × 256 × 1 (single slice), and resolution = 0.8 × 0.8 × 2 mm^3^. The multi-TI acquisition and short TE were designed to capture short-T1 components for subsequent spectral analysis.

#### 2.2.2 MRI of WMH tissue blocks on a preclinical 3T system

To establish an imaging strategy bridging postmortem hemisphere MRI and histopathology, tissue specimens were dissected into blocks under guidance from hemisphere MRI (**Fig. S1**). MRI experiments were performed on a preclinical 3T MRI system (BioSpec, Bruker BioSpin, Billerica, MA, USA) equipped with actively shielded gradients (maximum gradient strength = 900 mT/m; slew rate = 4200 T/m/s). We used a 40-mm inner-diameter transmit-receive quadrature coil with the following MRI protocols:

(**a**) **2D T2-FLAIR**: TI/TE/TR = 1100/69/4000 ms; matrix size = 200 × 200 × 1; resolution = 0.2 × 0.2 × 1.5 mm^3^.
(**b**) **2D T2-weighted Rapid Acquisition with Relaxation Enhancement (RARE)**: TE/TR = 80/3000 ms, RARE factor = 8, matrix size = 200 × 200 × 1, resolution = 0.2 × 0.2× 1.5 mm^3^.
(**c**) **Short-T1 MRI**: We used an inversion-recovery–prepared RARE (IR-RARE) sequence with 19 Tis, TE/TR same as in section 2.2.1d, RARE factor = 16, matrix size = 200 × 200 × 1, and resolution = 0.2 × 0.2 × 1.5 mm^3^.
(**d**) **T1-T2 correlation MRI:** To examine the relationship between short-T1 and short-T2 relaxation components, two-dimensional T1-T2 correlation MRI was performed. Nineteen inversion times (TIs), matching those used for the short-T1 MRI acquisition, were followed by a multi-echo readout (50 echo times; first TE = 6 ms; echo spacing = 6 ms, TR = 8000 ms), resulting in a 19 × 50 T1-T2 correlation dataset with matrix size = 100 × 100 × 1, and resolution = 0.4 × 0.4 × 1.5 mm^3^.

### 2.3 *In vivo* short T1 MRI

To assess the translational feasibility of the short-T1 fraction, short-T1 MRI was acquired in two living subjects: a 44-year-old healthy male individual without apparent neurological disorders and a 71-year-old female subject with a clinical diagnosis of moderate white matter microangiopathy, presenting WMHs lesion on T2-FLAIR images. All protocols were approved by the Institutional Review Board at New York University (NYU) Grossman School of Medicine. All participants provided written, informed consent before undergoing MRI scans.

The in vivo imaging protocol was analogous to the postmortem hemisphere specimen protocol described in section 2.2.1d, with the same TIs except that the 1,600 ms TI was replaced with 1,800 ms TI, effective TE/TR = 8.5/8000 ms, turbo factor = 4, matrix = 144 × 144 × 1, and voxel size = 1.5 × 1.5 × 3 mm^3^. To visualize the WMHs lesions, standard clinical 2D axial FLAIR and axial T2 weighted images were acquired with matching spatial resolution.

### 2.4 Image data processing

The T1-weighted signal within a selected voxel was modeled as a weighted integral over subvoxel components with distinct T1 relaxation times:

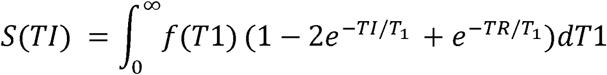

Signals acquired at multiple TIs signals were first polarity-corrected by inverting the data before the zero-crossing point and then analyzed using an inverse Laplace transform (ILT) with singular value decomposition and iterative Butler–Reed–Dawson optimization. A regularization parameter of 10^-6^ was used to estimate the distribution of T1 values, *f*(T1), within the range of TIs used and were calculated by integrating over all the peak area and further divided by the total peak area.

Based on previous reports of myelin water T1 relaxation times (18), the short-T1 fraction was defined as the fraction of spins with T1 < 300ms, which are potentially associated with myelin water. The long-T1 fraction was defined as the fraction of spins with T1 values between 300 and 800ms, which potentially include intracellular and extracellular water (EC/IC). The fraction of spins with T1 values above 1500ms was attributed to CSF or free water. Voxel-wise fraction maps were generated to visualize the spatial distribution of the three components.

Using an extension of the one-dimensional ILT framework, joint T1-T2 correlation data were analyzed with a two-dimensional ILT to estimate the voxel-wise joint relaxation time distribution *f*(T1,T2), as previously described in several studies (20,34). This multidimensional decomposition approach enables the identification of potential coupling between short-T1 and short-T2 signal components. The raw joint T1-T2 signals before the zero-crossing points were polarity-corrected. 2D-ILT were then performed on the corrected T1-T2 signal as described by Venkataramanan *et al.* (35).

### 2.5 Histopathological quantification

Following MRI, five tissue blocks containing normal-appearing white matter (NAWM) and periventricular and subcortical WMHs were processed for histopathology on an automated tissue processor (ASP 6025, Leica Biosystems), paraffin-embedded, and sectioned at 8 µm thickness. For routine histological evaluation, sections were stained with Luxol Fast Blue (LFB) to assess myelin content and counterstained with hematoxylin and eosin (H&E) to delineate overall tissue architecture. LFB/H&E staining was performed by the NYU Experimental Pathology Core.

For quantitative analysis of microscopy data, stain-specific optical density was computed from LFB images by subtracting the red channel from the blue channel in Matlab. MBP-stained images were grayscale-normalized (0–255) prior to analysis. Optical density values for both LFB and MBP staining were computed using previously described methods (36). To evaluate the consistency of staining across samples, we quantified the optical density distributions of LFB and MBP within the white matter regions of the three samples. The corresponding optical density histograms are presented in **Fig. S2**.

### 2.6 Statistical analysis

For each tissue specimen, four regions of interest (ROIs) encompassing WMHs and NAWM were manually delineated on T2-FLAIR images, and mean short-T1 fractions were extracted within each ROI. Matching ROIs were delineated on LFB- and MBP-stained histological sections, and mean optical density (OD) values was recorded. Pearson correlation analyses between short-T1 fractions and OD values were performed using Prism (GraphPad).

## 3. Results

### 3.1 WMHs in postmortem human brains had reduced short T1 components

Multiple WMH lesions were detected on T2-FLAIR images as hyperintense regions (**Fig. 1A**). In mTI images (**Fig. 1B**), the contrast between WMH and neighboring white matter varied as a function of TI. The corresponding quantitative T1 map (**Fig. 1C**) showed increased T1 values in WMH relative to NAWM. Voxel-wise short- and long-T1 fraction maps derived from ILT analysis (**Fig. 1E-F**) revealed a reduction in the short-T1 water fraction within WMH and a corresponding increase in the long-T1 fraction. Representative inversion-recovery signal curves from a WMH lesion (**Fig. 1F**) exhibited slower recovery than those from NAWM (**Fig. 1G**). The corresponding ILT results (**Fig. 1H & I**) showed a reduced short-T1 water peak and a shift toward longer T1 values in WMH, indicating a loss of short-T1 components.

**Figure 1.**
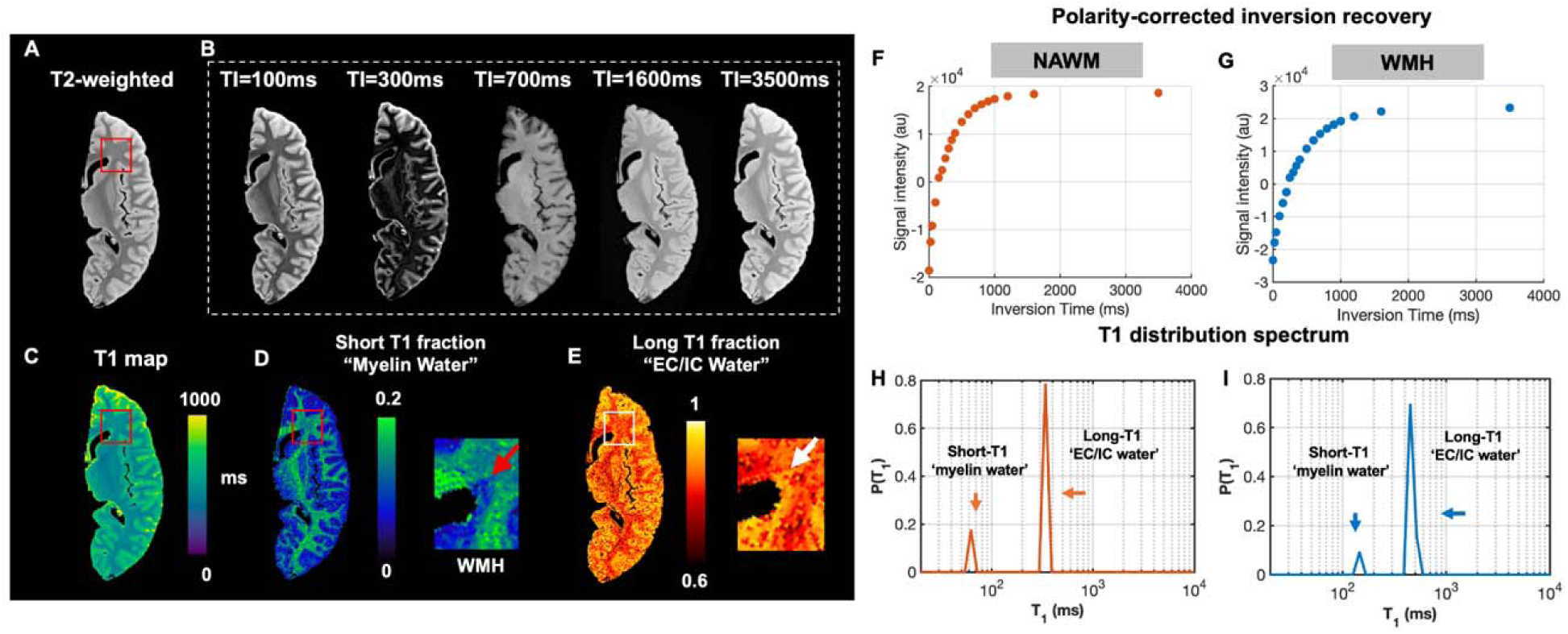
Postmortem brain hemisphere MRI. (**A**) Representative T2-FLAIR image identifying a periventricular white matter hyperintensity (WMH). (**B-C**) Multiple inversion recovery images acquired and the corresponding T1 map. (**D–E**) Voxel-wise shoft-T1 and long-T1 maps demonstrate reduced short-T1 fractions and elevated long-T1 fractions within WMHs. (**F–G**) Representative signal intensity profiles from normal-appearing white matter (NAWM) and WMH after polarity correction. (**H–I**) Corresponding ILT-derived T1 spectra showing discrete short- and long-T1 components in NAWM (**H**) and WMH (**I**), with a relative reduction of the short-T1 component in WMH.

### 3.2 High-resolution tissue block imaging revealed heterogeneous short-T1 contrast within WMHs

T2-weighted images of tissue blocks reproduced the hyperintense appearance of WMH lesions observed at the hemispheric level (**Fig. 2A-B**). At an in-plane resolution of 0.2 x 0.2 mm^2^, substantial intralesional heterogeneity became apparent on T2-weighted images. Short-T1 fraction maps further revealed a lesion core with short-T1 fractions approaching zero, surrounded by regions with intermediate short-T1 fractions relative to NAWM. Long-T1 fractions were increased in the lesion core compared with NAWM. In tissue blocks without apparent WMH (**Fig. 2C**), short-T1 fractions were relatively homogenous (∼ 0.1 -0.2).

**Figure 2.**
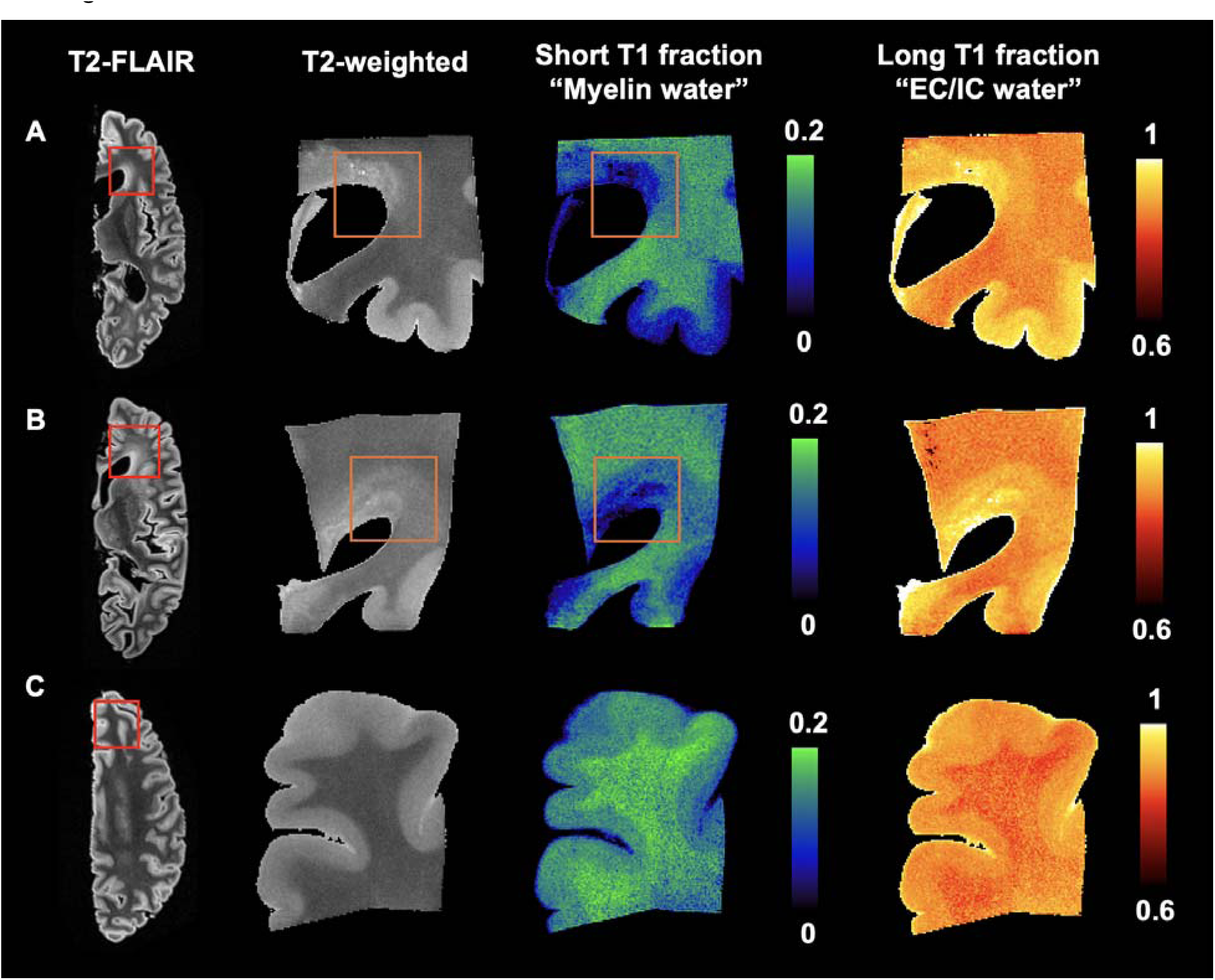
Postmortem tissue block MRI. (**A-B**) Images from tissue blocks containing periventricular WMHs. From left to right: hemispheric images showing periventricular WMHs in two specimens, T2-weighted images of the corresponding tissue blocks, and short- and long-T1 fraction maps. (**C**) Images from a tissue block containing NAWM with relatively uniform short-T1 fractions.

### 3.3 Relationship between short-T1 and short-T2 components characterized by T1-T2 correlation MRI

T1-T2 correlation analysis of NAWM and WMH revealed two distinct peaks (**Fig. 3A–B**): one characterized by shorter T1 and T2 values, and another by longer T1 and T2 values. Notably, the short-T1 and short-T2 components co-localized to the same peak in the T1-T2 spectrum, suggesting that these components arise from the same water pool, likely corresponding myelin-associated water. The separation between the two peaks along the T2 dimension (**Fig. 3C–D**) was markedly narrower than the separation along the T1 dimension (**Fig. 1H–I**), indicating that T1 provides greater contrast between the two water pools than T2 under the present acquisition conditions.

**Figure 3.**
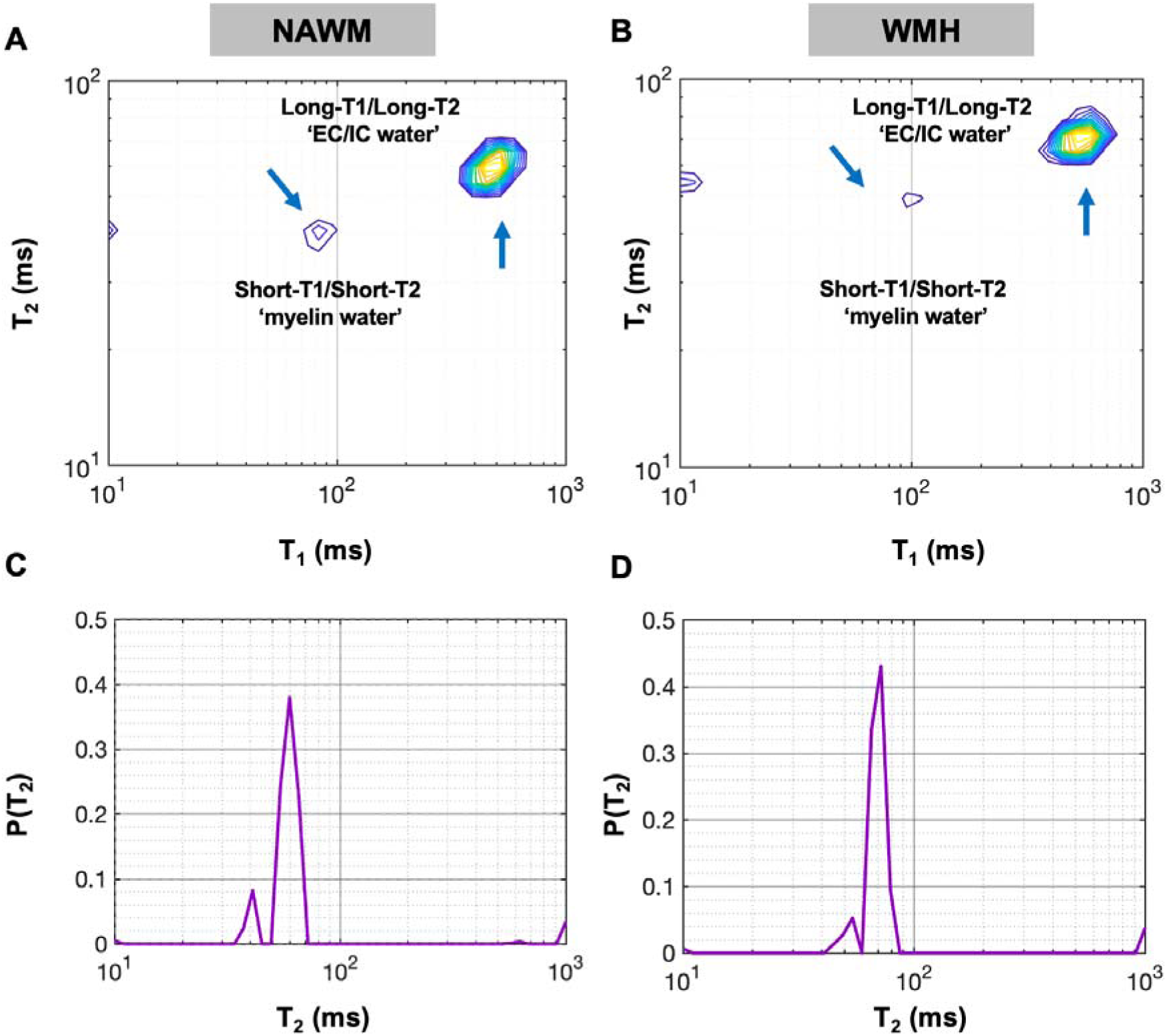
T1-T2 correlation spectra in NAWM (**A**) and WMH (**B**) and corresponding 1D projections along the T2 dimension (**C-D**).

### 3.4 Correlations between short-T1 fraction and myelin pathology

Matched histological staining and tissue block MRI were obtained from five tissue blocks. At low magnification, WMHs lesions in the tissue blocks appeared pale compared with NAWM on LFB-stained sections (**Fig. 4A, C**), whereas MBP-stained sections (**Fig. 4B, D**), showed only a mild reduction in staining intensity. At higher magnification, loss of myelinated fibers within WMH became evident on both LFB- and MBP-stained sections. In MBP-stained WHM regions, more pronounced vacuolation and rarefaction of the WM were observed compared with NAWM, consistent with myelin degeneration and tissue rarefaction.

**Figure 4.**
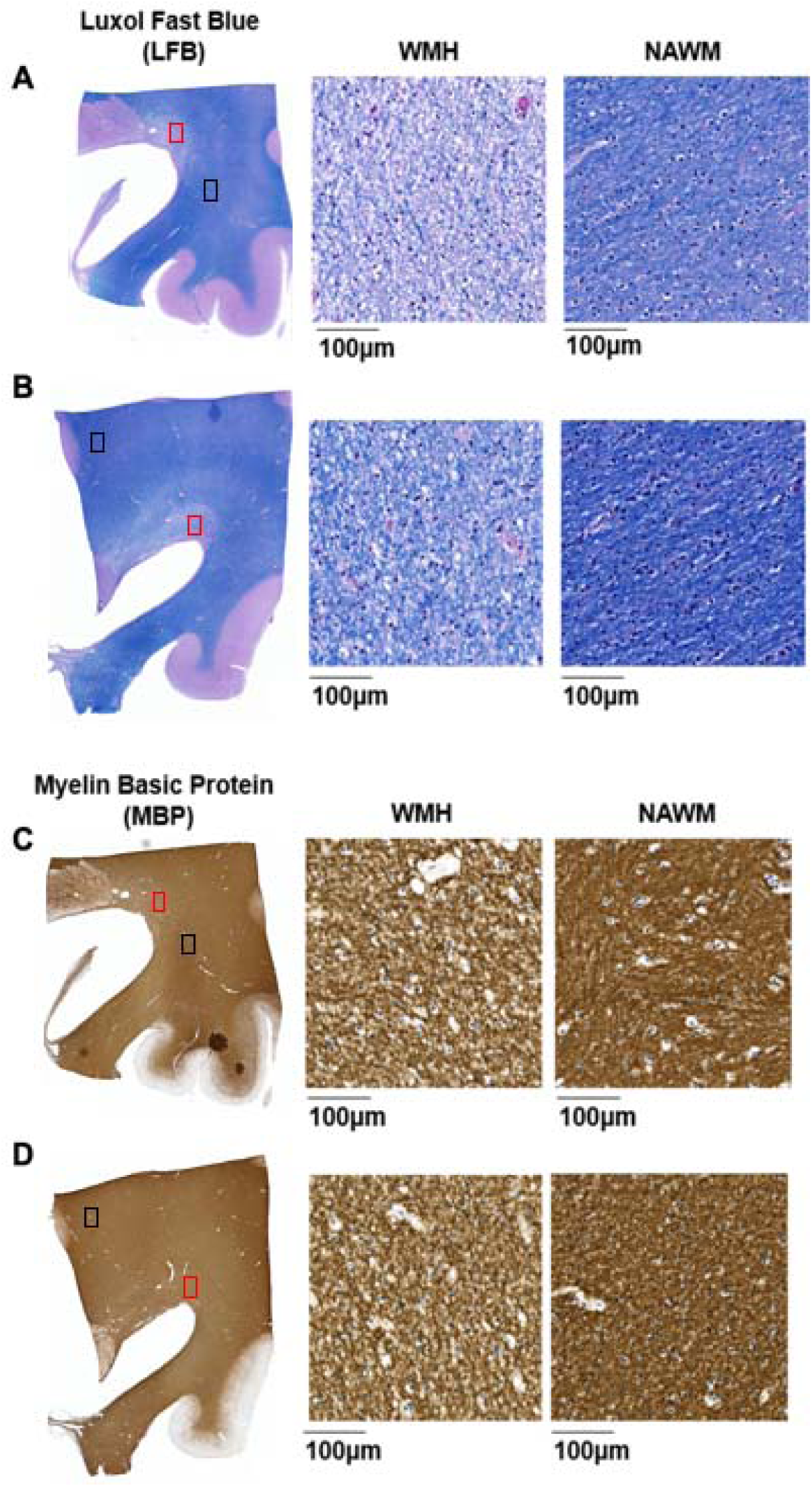
Luxol Fast Blue (LFB) and myelin basic protein (MBP) stained histological sections from two tissue blocks containing white matter hyperintensities (WMHs). (A–B) LFB-stained sections and corresponding high-magnification images of WMH and normal-appearing white matter (NAWM). (C–D) MBP-stained sections and corresponding high-magnification images of WMH and NAWM.

Regions of interest (ROIs) were manually delineated on both MRI and histological sections at matching spatial locations (**Fig. 5A–F**). Four ROIs within white matter were defined for each sample, yielding a total of 20 ROIs for MRI–histopathological correlation (**Fig. S3**). Short-T1 fractions were significantly correlated with histological measures of myelin content across NAWM and WMH ROIs. Specifically, short-T1 fraction showed a strong sigmoidal relationship with LFB OD (R² = 0.829, **Fig. 5G**) and a robust linear relationship with MBP OD (R² = 0.685, **Fig. 5H**), indicating that reductions in short-T1 fraction are tightly linked to myelin loss in WMHs.

**Figure 5.**
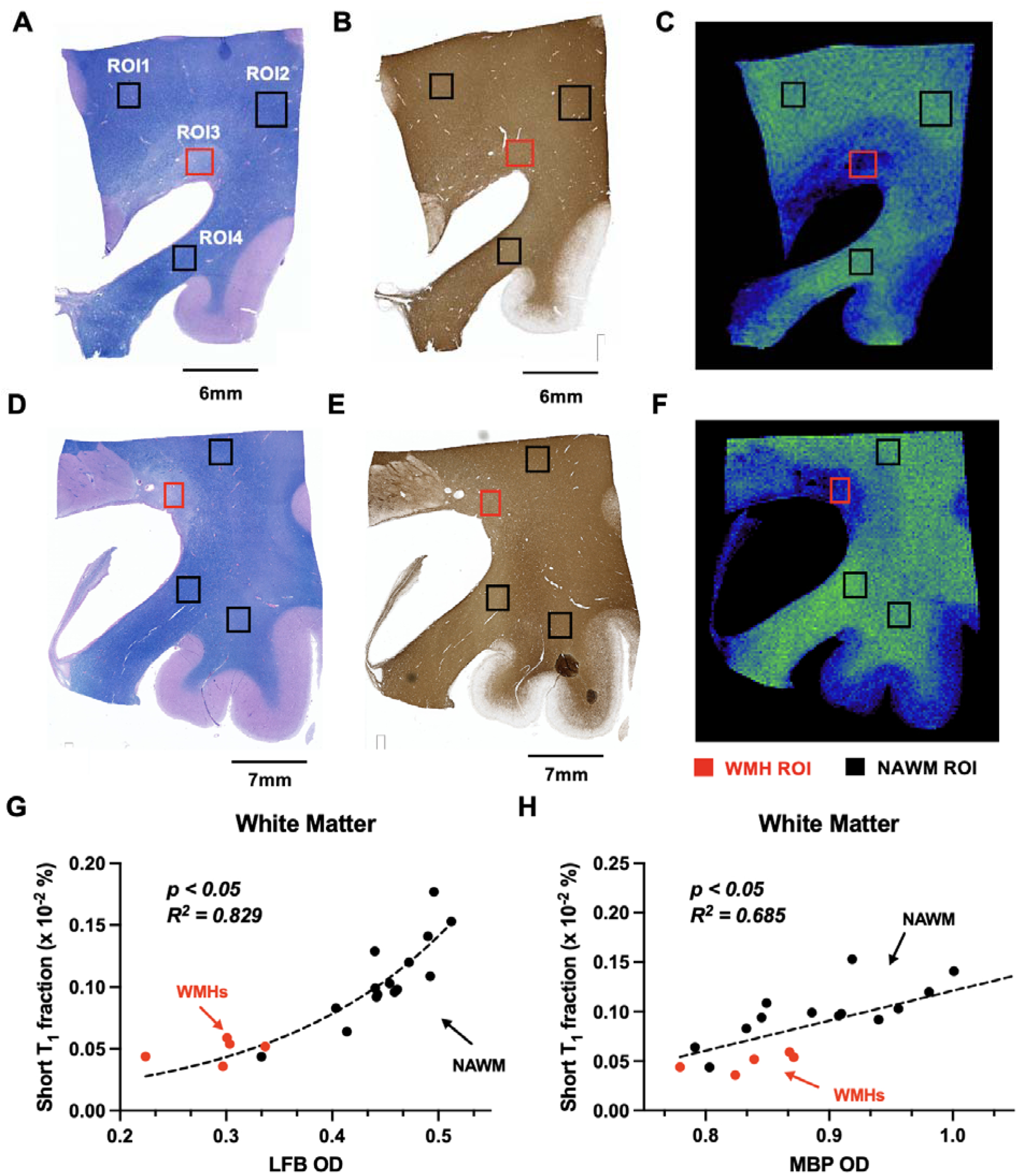
Correlation of short-T1 fraction with histological measures of myelin content. (A, D) Luxol Fast Blue (LFB)-stained sections. (B, E) Myelin basic protein (MBP)-stained sections. (C, F) Corresponding short-T1 fraction maps derived from MRI. Red and black rectangles indicate white matter hyperintensity (WMH) and normal-appearing white matter (NAWM) regions of interest (ROIs), respectively. (G) Correlation between short-T1 fraction and LFB optical density (OD) across white matter ROIs. (H) Correlation between short-T1 fraction and MBP optical density across white matter ROIs.

### 3.5 *In vivo* mapping showed reduced short-T1 fraction of WMHs

WMHs were identified primarily on clinical FLAIR and T2-weighted MRI (**Fig. 6A-B**). Consistent with the postmortem findings, short-T1 fraction maps demonstrated reduced short-T1 fractions within all three WMH regions in one subject (**Fig. 6C**). In one WMH region, the short-T1 reduction was accompanied by increased long-T1 fractions (**Fig. 6D**), with no obvious increase in CSF/free-water fraction (**Fig. 6E**), but the other two WMHs show no obvious increase in long-T1 fractions, potentially reflecting different stages and types of WMH pathology.

**Figure 6.**
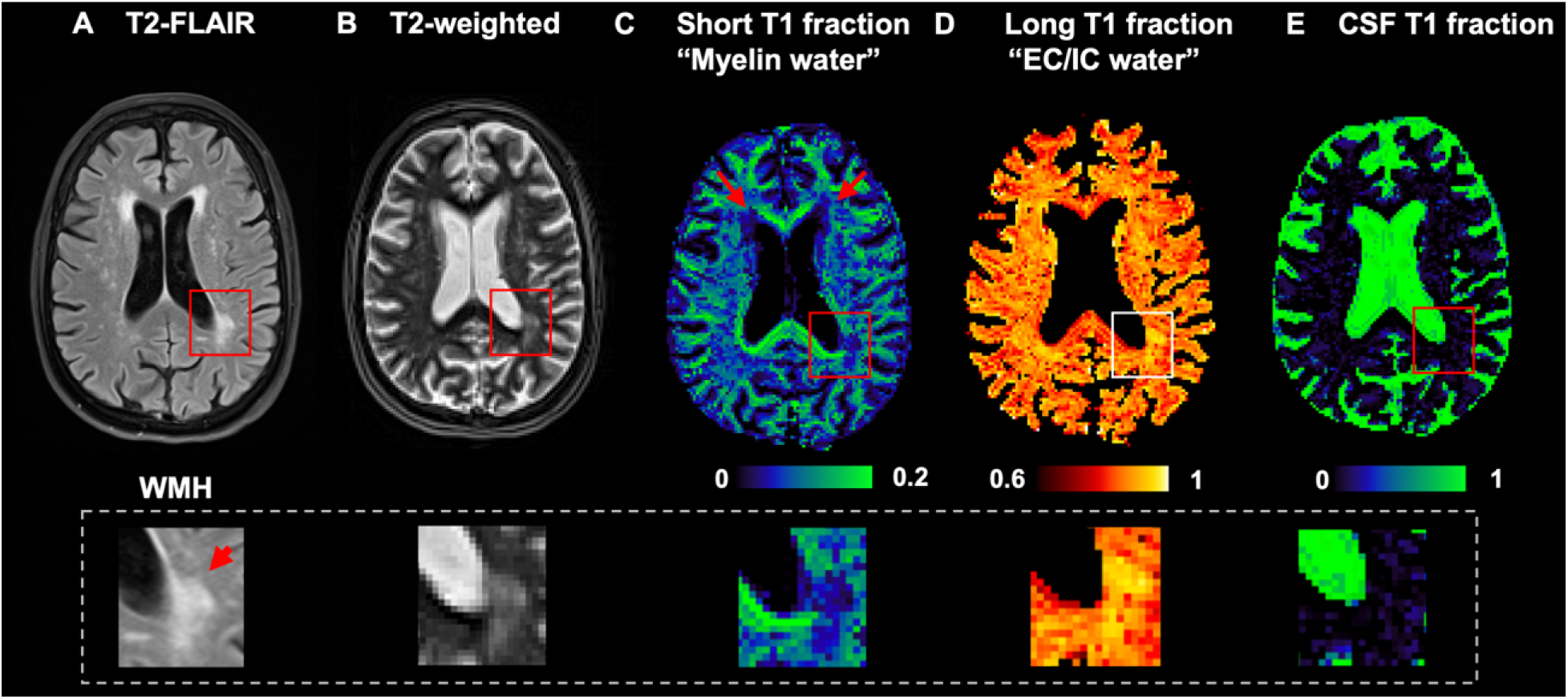
*In vivo* short-T1 fraction mapping in a patient with WMHs. (**A–E**) Patient with a periventricular WMH showing T2-FLAIR (**A**), T2-weighted imaging (**B**), and corresponding short-T1 (putative myelin water; **C**), long-T1 (EC/IC water; **D**), and CSF/free-water T1 fraction maps (**E**). Reduced short-T1 fraction is observed within the WMH region (red/white box).

## 4. Discussion

In this study, we demonstrated that the short-T1 fraction derived from multi-inversion-time T1 relaxometry is sensitive to myelin-related alterations in WMHs. The short-T1 fractions in WMHs and NAWM strongly correlated with myelin-stained histology. *In vivo* results also demonstrate similar reductions in short-T1 fractions in WMHs, supporting the translational potential of this approach.

### 4.1 Short-T1 fraction as an alternative approach for myelin water imaging

Over the past several decades, MRI has been widely used to assess myelin and MWI has emerged as a powerful technique to quantify myelin integrity. Conventional MWI is most often performed using multi-component T2 analysis (8). The contrast mechanism relies on the unique MR properties of water located near the lipid bilayers of the myelin sheath, which experience restricted motion and strong macromolecular interactions, leading to markedly shorter T1 and T2 values than those of intra- and extracellular water compartments. In T2-based MWI, multi-echo spin-echo or gradient and spin echo (GRASE) acquisitions(37–40) are used to record signal decay curves that can be decomposed into distinct T2 components using approaches such as ILT or non-negative least squares (NNLS)(41,42).

Conventional T2-based MWI typically requires long echo trains with high signal-to-noise ratio (SNR) and dense echo-time sampling, which can increase scan time and make the method more sensitive to B1 inhomogeneity and stimulated echo contamination (19). Methodological advances have included the use of GRASE to accelerate acquisition and improve spatial coverage (37), as well as improved postprocessing strategies for more accurate short-T2 fraction estimation (43). Importantly, several validation studies have demonstrated strong correlations between T2-derived myelin water fraction and histological measures of myelin in multiple sclerosis and other myelin pathologies (10,44).

In parallel with T2-based MWI, T1 relaxation characteristics of myelin water have also been investigated as complementary markers of myelin microstructure. Labadie *et al.* (18) demonstrated that, analogous to multi-echo T2 relaxometry, multicomponent T1 relaxation can be resolved using a multiple inversion-recovery sampling strategy to capture multiexponential longitudinal recovery. Using ILT, they identified distinct T1 components within brain tissue, including a prominent short-T1 fraction predominantly localized to white matter and interpreted as reflecting myelin-associated water. Our T1-T2 correlation results further support this interpretation. The concordance between short-T1 and short-T2 components (**Fig. 3**) indicates that both parameters are probing the same underlying water pool. Only a limited number of studies have directly examined short-T1 and short-T2 relaxation components using joint T1-T2 correlation spectroscopy in myelinated tissue (20,45–47), and evidence from human postmortem white matter at 3T remains sparse. Collectively, these studies suggest that myelin water exhibits both shortened T1 and T2 values.

Compared with T2-based MWI, T1-based approaches are intrinsically less sensitive to B1 inhomogeneity and are not affected by stimulated echo contamination. Furthermore, as demonstrated by the T1-T2 correlation results (**Fig. 3A-B**), the short-T1 component exhibits greater spectral separation from longer-T1 components (on the order of 10^1^-10^3^ ms) than the separation between short- and long-T2 components (10^1^ - 10^2^). This suggests that T1 provides greater discriminatory power than T2 for separating myelin water from other tissue water pools in postmortem white matter under our acquisition conditions. A practical implication is that the short-T1 compartment can be robustly resolved using a relatively broad and potentially sparser inversion-time sampling scheme, whereas accurate separation of short-T2 components may depend more critically on dense echo-time sampling and higher SNR. This may also represent a potential advantage for MWI at high magnetic field strengths (7T or higher), as the distance between short-T2 and long-T2 components may be reduced due to the overall shortening of T2 values at higher fields, while T1 generally increases with field strength.

### 4.2 Short T1 Myelin Water Imaging for assessing subvoxel myelin injury in WMHs

In this study, we implemented short-T1 MRI on both clinical and preclinical MRI 3T systems to investigate the short-T1 myelin water fraction in postmortem WM containing WMHs. WMHs are frequently observed on T2-weighted and T2-FLAIR images in older adults and in AD patients. Their pathophysiology is multifactorial (48), involving chronic hypoperfusion (49), impaired vascular reactivity (50), demyelination, reactive gliosis, and WM rarefaction with increased extracellular water (51,52). Among these processes, demyelination is increasingly recognized as a central contributor to white matter injury and may represent an early pathological event preceding overt tissue loss. Sensitive, noninvasive imaging biomarkers that can detect subtle myelin alterations before macroscopic WMHs become apparent would therefore be highly valuable for elucidating disease mechanisms and facilitating early intervention strategies.

In our postmortem hemispheric MRI scans of three AD brains with WMH lesions, the short-T1 fraction map exhibited strong contrast between white matter and cortex (**Fig. 1D**), consistent with the higher myelin content of white matter. Within WMH regions, the short-T1 fraction was markedly reduced relative to surrounding NAWM, indicating that the short-T1 compartment is sensitive to myelin loss associated with WMH pathology. These findings align with prior histopathological reports that WMHs are characterized by demyelination, axonal loss, gliosis, and WM rarefaction with increased extracellular water content (51,53). Such microstructural alterations are expected to reduce the relative contribution of short-T1 components (myelin-associated water) and increase the relative contribution of longer-T1 pools originating from intra- and extracellular water. The observed decrease in short-T1 fraction within WMHs lesions is therefore consistent with the underlying pathological features of white matter degeneration.

We further performed short-T1 fraction mapping in tissue blocks on a preclinical 3T system, which enabled substantially higher spatial resolution and close MRI-histopathology correspondence. At high spatial resolution, heterogeneous patterns of short-T1 fraction reduction were observed within WMH lesions, with more pronounced decreases in the lesion core and relatively milder reductions in surrounding penumbral regions (**Fig. 2A-B**). These spatial variations likely reflect different stages or severities of myelin degeneration within the lesion. Such patterns are consistent with neuropathological studies demonstrating that WMHs often exhibit a gradient of injury severity, ranging from severe demyelination and axonal loss in the core to partial myelin disruption and WM rarefaction in the periphery. The ability of short-T1 fraction imaging to capture these spatially varying alterations suggests that this approach may provide a sensitive marker for assessing the degree and heterogeneity of myelin pathology within WM disease.

To further evaluate short-T1 fraction as a myelin water biomarker, we performed correlation analyses between short-T1 fraction and LFB and MBP-stained histology. LFB provides a robust measure of myelin content, whereas MBP immunostaining offers greater molecular specificity for myelin sheaths. Both markers confirmed myelin loss, including both myelin pallor and increased vacuolation (**Fig. 4**), within WMHs in postmortem tissue. Short-T1 fraction showed strong correlations with both LFB and MBP optical density, with R^2^ comparable to previous reports on correlations between T2-based myelin water fractions and myelin histology in different pathological contexts (10,44). For example, Laule et al.(44) reported an average R^2^ of 0.78 between short-T2 fraction and LFB staining in formalin fixed human brain specimens with multiple sclerosis. To minimize potential confounding from gray-white matter differences in microstructure, all correlations were restricted to white matter ROIs in this study.

Interestingly, the relationship between short-T1 fraction and MBP OD was approximately linear, whereas the relationship with LFB staining was better described by a sigmoidal function. In regions of severe myelin loss, short-T1 fraction increased slowly with rising LFB signal, whereas in NAWM the relationship was much steeper. One potential explanation is that the T1-shortening effect of myelin on nearby water is not strictly proportional to myelin content, particularly at low myelin densities, leading to a nonlinear relationship between myelin content and short-T1 fraction. LFB and MBP target different myelin components and may differ in their sensitivity and dynamic range across the spectrum of myelin loss, and OD is a semi-quantitative measurement. As a result, one marker may attenuate faster than the other as myelin decreases. Future work incorporating more detailed biophysical modeling and quantitative myelin markers (e.g. G-ratio from electron microscopy) will be important to clarify these relationships.

### 4.3 Feasibility of measuring short-T1 myelin water fraction *in vivo*

Our preliminary *in vivo* results demonstrate the feasibility of translating short-T1 fraction imaging to living human subjects. As shown in **Fig. 6A-B**, conventional T2-FLAIR and T2-weighted images revealed a representative periventricular WMH. The corresponding short-T1 fraction map exhibited spatial patterns consistent with known myelin distributions (e.g. elevated short T1 fraction in corpus callosum) and showed a clear reduction within the periventricular WMH (**Fig. 6C**). Conversely, the long-T1 fraction, attributed to intra- and extracellular water compartments, was relatively increased within the WMH lesion (**Fig. 6D**), consistent with increased extracellular water content associated with white matter degeneration.

In addition, a free-water pool with very long T1 relaxation times (T1 > 3000 ms) was isolated to represent CSF-like signal (**Fig. 6E**). This component showed elevated contribution near the ventricular boundaries and, to a lesser extent, within periventricular WMHs, suggesting increased free-water content, edema, and/or partial CSF contamination in these regions. Together, these results indicate that T1 fraction mapping can separate distinct water compartments *in vivo* and may provide complementary microstructural information beyond conventional structural MRI.

It is important to recognize that absolute T1 values differ between *ex vivo* and *in vivo* tissue due to factors such as fixation, temperature, and postmortem biochemical changes (33). Nonetheless, the overall spatial distribution of the *in vivo* short-T1 fraction maps remained consistent with known patterns of myelination in the human brain, supporting the biological plausibility of the metric.

### 4.4 Technical considerations and limitations

Several technical considerations merit discussion. First, the short-T1 fraction is determined by longitudinal magnetization dynamics and, in principle, can be estimated using different readout strategies. In this study, we used a TSE readout to minimize sensitivity to B₀ inhomogeneity and T2*-related susceptibility effects. However, sufficiently short effective echo times are required to limit additional T2- and T2*-dependent signal loss from the intrinsically short-T2 myelin water pool. Excessive T2/T2* weighting during the readout does not alter the underlying T1 recovery itself, but it can disproportionately suppress the contribution of short-T2 components to the measured signals, thereby biasing the estimated short-T1 fraction. The influence of echo time on short-T1 fraction estimates was not systematically evaluated here due to scan time constraints and warrant further investigation.

Second, we employed a single-slice acquisition to demonstrate feasibility of short-T1 quantification. In conventional 2D multi-slice acquisitions, slice-timing differences across inversion recovery periods can introduce inconsistencies in the effective T1 for each slice. More advanced sampling strategies, such as slice-shuffling (54,55) approaches, may help mitigate these effects and enable accelerated multi-slice acquisition. 3D acquisitions with nonselective inversion pulses can provide whole-brain coverage for short-T1 fraction mapping but generally require longer scan times than 2D implementations. Optimization of sequence parameters and integration of acceleration techniques (e.g., parallel imaging, compressed sensing) (56,57) will be important to balance spatial coverage, spatial resolution, and scan efficiency in clinical applications.

Third, our postmortem MRI was performed at 3T using both clinical and preclinical systems, facilitating clinical translation at the same field strength. Although several myelin-sensitive MRI techniques have been evaluated across different field strengths (58,59), further investigation is needed to determine optimal short-T1 spectral selection at higher fields, given that T1 values tend to increase with field strength and T2 values tend to decrease with field strength. The advantage of the short-T1 may alter the separation between compartments.

This study has several limitations. First, the translational interpretation of short-T1 fractions is constrained by altered relaxation properties in postmortem tissue. Factors such as formalin fixation (60–62), tissue rehydration (63), and variability in postmortem interval (64–66) can all systematically influence T1 values and relative compartment fractions. Although specimens were processed using standardized ADRC protocols, these factors should be considered when extrapolating *ex vivo* findings to *in vivo* conditions. Second, the number of postmortem samples in this study was limited. Larger cohorts, including subjects with diverse WM pathologies and longitudinal *in vivo* follow-up, will be essential for validating and generalizing the current findings. Third, although both periventricular and subcortical WMHs lesions were included in this study, a systematic comparison between these two lesion types was not performed due to the limited sample size. Future studies with larger number of brain specimens with WMHs would enable a more comprehensive evaluation of the pathological differences between periventricular and subcortical WMHs, which may further help differentiate these lesion types using short-T1 imaging measures(67). Lastly, although Labadie et al.(18) discussed the potential effects of myelin water exchange between short-T1 myelin-associated water and other tissue water pools on the observed short-T1 peak distribution, such effects were not systematically evaluated in this study. Increased myelin water exchange may occur in WMH lesions with degraded myelin sheaths, potentially shifting the system toward a faster exchange regime and thereby reducing and broadening the observable short-T1 fraction. In addition, the estimated short-T1 peak distribution may also depend on the inversion recovery sampling scheme, particularly the maximum inversion time, which can influence the apparent position and fraction of the short-T1 component. Further investigation of exchange effects and acquisition-dependent biases may therefore provide additional insight into the biophysical interpretation of short-T1 measurements and their relationship to myelin water exchange.

## 5. Conclusion

In conclusion, subvoxel short-T1 water fraction correlated strongly with LFB and MBP-stained myelin histology in postmortem human white matter and WMHs, supporting its validity as a quantitative marker of myelin. The *ex vivo* and initial *in vivo* findings highlight the feasibility of using short-T1 fraction imaging as a myelin-sensitive biomarker for assessing white matter integrity in WMHs.

## Data Availability Statement

The data that support the findings of this study are available on request from the corresponding author. The data is not publicly available due to privacy or ethical restrictions.

---

**Fig. S1.**
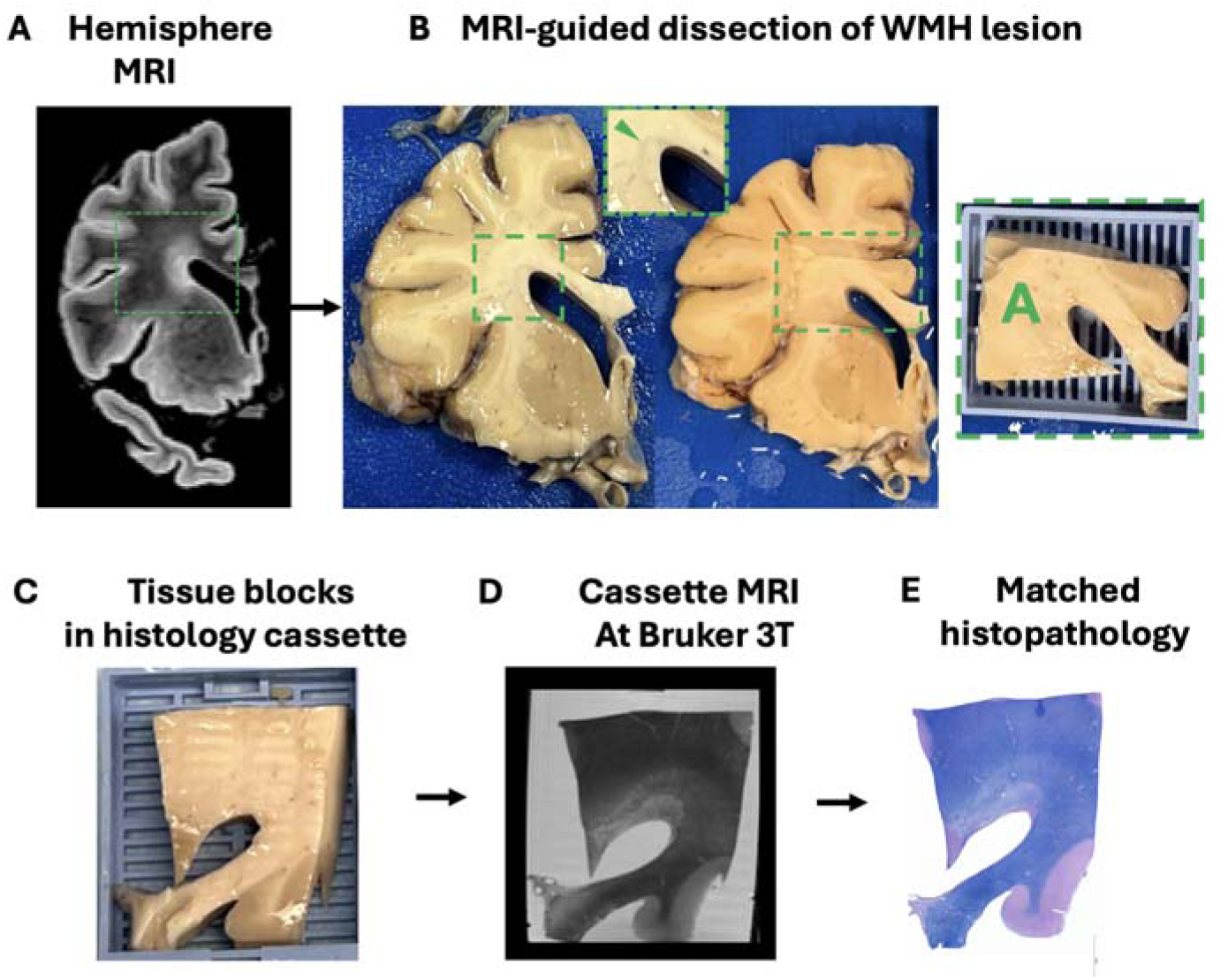
Representative MRI-guided dissection and MRI–histology matching of white matter hyperintensity (WMH) tissue. **(A-B)** *Ex vivo* hemisphere MRI was used to localize the WMH lesion and guide targeted tissue dissection. **(C-D)**The selected tissue block was placed in a histology cassette and imaged using high-resolution cassette MRI at 3T prior to processing. **(E)** Corresponding histological sections were subsequently obtained from the same block, enabling spatially matched MRI–histopathology comparison.

**Fig. S2.**
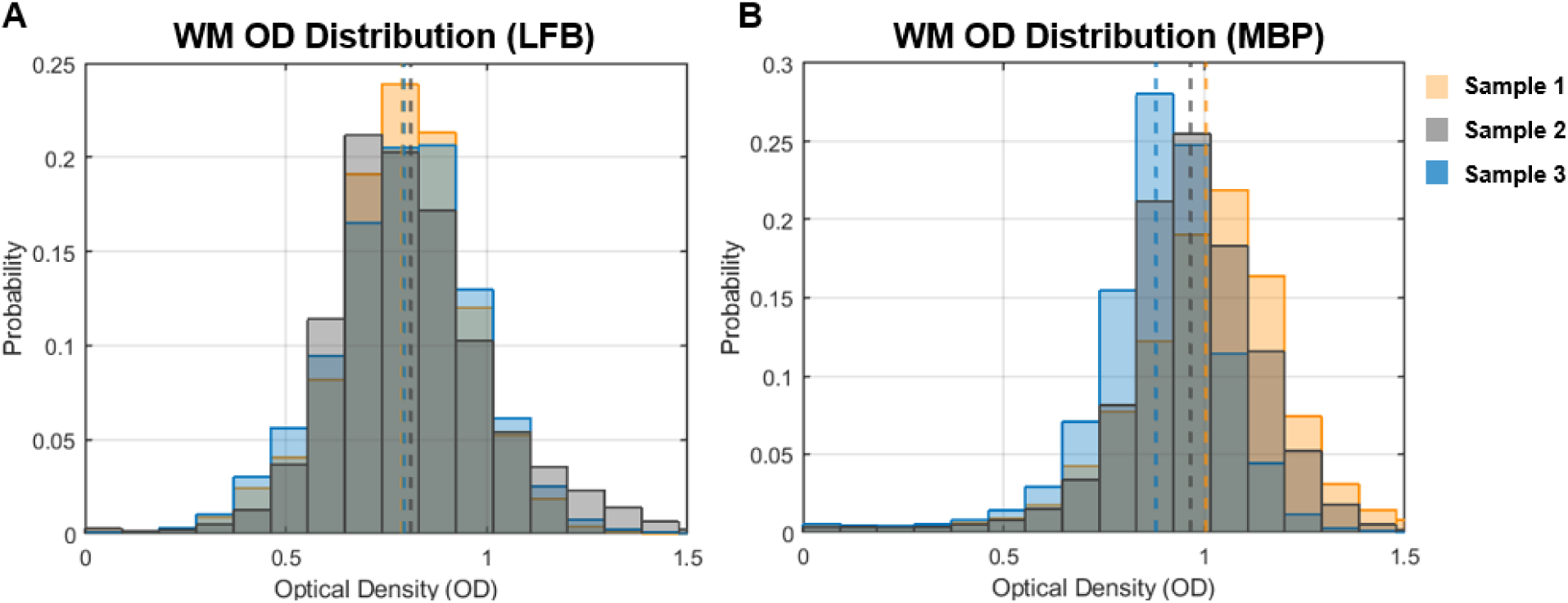
Distribution of normalized optical density (OD) values across three different samples for (A) Luxol Fast Blue (LFB) staining and (B) myelin basic protein (MBP) staining in white matter (WM). Dashed lines indicate the mean OD value for each sample.

**Fig. S3.**
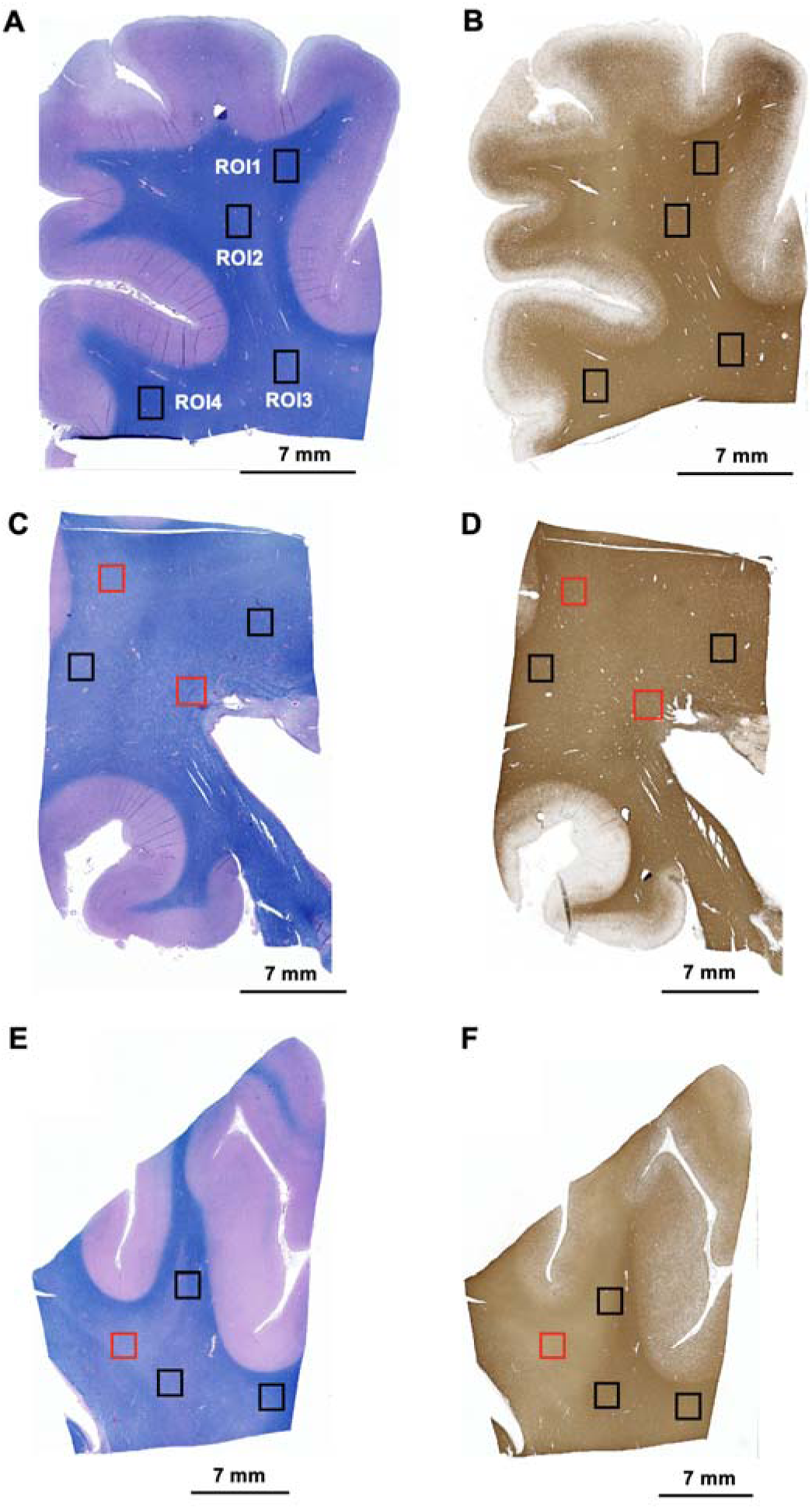
ROI placement on histology in additional samples. Representative examples of white matter ROI placement on Luxol Fast Blue (A, C, E) and myelin basic protein (B, D, F) stained sections from additional samples are shown. A total of 20 ROIs were included across all samples. The red squares in C-F indicate corresponding ROIs for WMHs lesions.

## Reference

1. Peter C, Sathe A, Shashikumar N, Pechman KR, Workmeister AW, Jackson TB, Huo Y, Mukherjee S, Mez J, Dumitrescu LC, Gifford KA, Bolton CJ, Gaynor LS, Risacher SL, Beason-Held LL, An Y, Arfanakis K, Erus G, Davatzikos C, Tosun-Turgut D, Habes M, Wang D, Toga AW, Thompson PM, Zhang P, Schilling KG, Albert M, Kukull W, Biber SA, Landman BA, Bendlin BB, Johnson SC, Schneider J, Barnes LL, Bennett DA, Jefferson AL, Resnick SM, Saykin AJ, Crane PK, Cuccaro ML, Hohman TJ, Archer DB, and the Alzheimer’s Disease Sequencing Project Phenotype Harmonization Consortium Analyst T, Zaras D, Yang Y, Durant A, Kanakaraj P, Kim ME, Gao C, Newlin NR, Ramadass K, Khairi NM, Li Z, Yao T, Choi SE, Klinedinst B, Lee ML, Scollard P, Trittschuh EH, Sanders EA. White Matter Abnormalities and Cognition in Aging and Alzheimer Disease. JAMA Neurol 2025;82(8):825–836.

2. Mito R, Raffelt D, Dhollander T, Vaughan DN, Tournier JD, Salvado O, Brodtmann A, Rowe CC, Villemagne VL, Connelly A. Fibre-specific white matter reductions in Alzheimer’s disease and mild cognitive impairment. Brain 2018;141(3):888–902.

3. Prins ND, Scheltens P. White matter hyperintensities, cognitive impairment and dementia: an update. Nat Rev Neurol 2015;11(3):157–165.

4. Alber J, Alladi S, Bae HJ, Barton DA, Beckett LA, Bell JM, Berman SE, Biessels GJ, Black SE, Bos I, Bowman GL, Brai E, Brickman AM, Callahan BL, Corriveau RA, Fossati S, Gottesman RF, Gustafson DR, Hachinski V, Hayden KM, Helman AM, Hughes TM, Isaacs JD, Jefferson AL, Johnson SC, Kapasi A, Kern S, Kwon JC, Kukolja J, Lee A, Lockhart SN, Murray A, Osborn KE, Power MC, Price BR, Rhodius-Meester HFM, Rondeau JA, Rosen AC, Rosene DL, Schneider JA, Scholtzova H, Shaaban CE, Silva N, Snyder HM, Swardfager W, Troen AM, van Veluw SJ, Vemuri P, Wallin A, Wellington C, Wilcock DM, Xie SX, Hainsworth AH. White matter hyperintensities in vascular contributions to cognitive impairment and dementia (VCID): Knowledge gaps and opportunities. Alzheimers Dement (N Y) 2019;5:107–117.

5. Groh J, Simons M. White matter aging and its impact on brain function. Neuron 2025;113(1):127–139.

6. Garnier-Crussard A, Bougacha S, Wirth M, Dautricourt S, Sherif S, Landeau B, Gonneaud J, De Flores R, de la Sayette V, Vivien D, Krolak-Salmon P, Chetelat G. White matter hyperintensity topography in Alzheimer’s disease and links to cognition. Alzheimers Dement 2022;18(3):422–433.

7. Nave KA, Werner HB. Myelination of the nervous system: mechanisms and functions. Annu Rev Cell Dev Biol 2014;30:503–533.

8. MacKay A, Whittall K, Adler J, Li D, Paty D, Graeb D. In vivo visualization of myelin water in brain by magnetic resonance. Magn Reson Med 1994;31(6):673–677.

9. Whittall KP, MacKay AL, Graeb DA, Nugent RA, Li DK, Paty DW. In vivo measurement of T2 distributions and water contents in normal human brain. Magn Reson Med 1997;37(1):34–43.

10. Laule C, Leung E, Lis DK, Traboulsee AL, Paty DW, MacKay AL, Moore GR. Myelin water imaging in multiple sclerosis: quantitative correlations with histopathology. Mult Scler 2006;12(6):747–753.

11. Alonso-Ortiz E, Levesque IR, Pike GB. Multi-gradient-echo myelin water fraction imaging: Comparison to the multi-echo-spin-echo technique. Magn Reson Med 2018;79(3):1439–1446.

12. Du YP, Chu R, Hwang D, Brown MS, Kleinschmidt-DeMasters BK, Singel D, Simon JH. Fast multislice mapping of the myelin water fraction using multicompartment analysis of T2* decay at 3T: a preliminary postmortem study. Magn Reson Med 2007;58(5):865–870.

13. van Gelderen P, de Zwart JA, Lee J, Sati P, Reich DS, Duyn JH. Nonexponential T(2) decay in white matter. Magn Reson Med 2012;67(1):110–117.

14. Lee J, Nam Y, Choi JY, Kim EY, Oh SH, Kim DH. Mechanisms of T(2) * anisotropy and gradient echo myelin water imaging. NMR Biomed 2017;30(4).

15. Lee J, Hyun JW, Lee J, Choi EJ, Shin HG, Min K, Nam Y, Kim HJ, Oh SH. So You Want to Image Myelin Using MRI: An Overview and Practical Guide for Myelin Water Imaging. J Magn Reson Imaging 2021;53(2):360–373.

16. Mancini M, Karakuzu A, Cohen-Adad J, Cercignani M, Nichols TE, Stikov N. An interactive meta-analysis of MRI biomarkers of myelin. Elife 2020;9.

17. van der Weijden CWJ, Biondetti E, Gutmann IW, Dijkstra H, McKerchar R, de Paula Faria D, de Vries EFJ, Meilof JF, Dierckx R, Prevost VH, Rauscher A. Quantitative myelin imaging with MRI and PET: an overview of techniques and their validation status. Brain 2023;146(4):1243–1266.

18. Labadie C, Lee JH, Rooney WD, Jarchow S, Aubert-Frecon M, Springer CS, Jr., Moller HE. Myelin water mapping by spatially regularized longitudinal relaxographic imaging at high magnetic fields. Magn Reson Med 2014;71(1):375–387.

19. Lebel RM, Wilman AH. Transverse relaxometry with stimulated echo compensation. Magn Reson Med 2010;64(4):1005–1014.

20. Travis AR, Does MD. Selective excitation of myelin water using inversion-recovery-based preparations. Magn Reson Med 2005;54(3):743–747.

21. Andrews TJ, Osborne MT, Does MD. Diffusion of myelin water. Magn Reson Med 2006;56(2):381–385.

22. Athertya JS, Daskareh M, Shin SH, Wang J, Lo J, Ma Y. Brain short T(2) component imaging using double adiabatic inversion recovery prepared ultrashort Echo time (DIR-UTE) sequence. Neuroimage 2025;316:121315.

23. Does MD. Relaxation-selective magnetization preparation based on T1 and T2. J Magn Reson 2005;172(2):306–311.

24. Oh SH, Bilello M, Schindler M, Markowitz CE, Detre JA, Lee J. Direct visualization of short transverse relaxation time component (ViSTa). Neuroimage 2013;83:485–492.

25. Oh SH, Lee G, Lee J. Three-Dimensional Efficient Myelin-Weighted Imaging Utilizing Direct Visualization of Short Transverse Relaxation Time Component (ViSTa). Hum Brain Mapp 2025;46(12):e70307.

26. Ma YJ, Jang H, Lombardi AF, Corey-Bloom J, Bydder GM. Myelin water imaging using a short-TR adiabatic inversion-recovery (STAIR) sequence. Magn Reson Med 2022;88(3):1156–1169.

27. Ma YJ, Jang H, Wei Z, Cai Z, Xue Y, Lee RR, Chang EY, Bydder GM, Corey-Bloom J, Du J. Myelin Imaging in Human Brain Using a Short Repetition Time Adiabatic Inversion Recovery Prepared Ultrashort Echo Time (STAIR-UTE) MRI Sequence in Multiple Sclerosis. Radiology 2020;297(2):392–404.

28. Moazamian D, Shaterian Mohammadi H, Athertya JS, Shin SH, Lo J, Chang EY, Du J, Bydder GM, Ma Y. Myelin water quantification in multiple sclerosis using short repetition time adiabatic inversion recovery prepared-fast spin echo (STAIR-FSE) imaging. Quant Imaging Med Surg 2024;14(2):1673–1685.

29. Shaterian Mohammadi H, Moazamian D, Athertya JS, Shin SH, Lo J, Suprana A, Malhi BS, Ma Y. Quantitative myelin water imaging using short TR adiabatic inversion recovery prepared echo-planar imaging (STAIR-EPI) sequence. Front Radiol 2023;3:1263491.

30. Deoni SC, Rutt BK, Arun T, Pierpaoli C, Jones DK. Gleaning multicomponent T1 and T2 information from steady-state imaging data. Magn Reson Med 2008;60(6):1372–1387.

31. Deoni SC, Kolind SH. Investigating the stability of mcDESPOT myelin water fraction values derived using a stochastic region contraction approach. Magn Reson Med 2015;73(1):161–169.

32. Nguyen-Hao HT, Liu J, Novelli M, Maskey D, Schwartz RS, Harari O, Sutherland GT. Quantification of Alzheimer disease neuropathology using tissue microarrays. J Neuropathol Exp Neurol 2025;84(10):855–869.

33. Shepherd TM, Thelwall PE, Stanisz GJ, Blackband SJ. Aldehyde fixative solutions alter the water relaxation and diffusion properties of nervous tissue. Magn Reson Med 2009;62(1):26–34.

34. Song YQ, Venkataramanan L, Hurlimann MD, Flaum M, Frulla P, Straley C. T(1)--T(2) correlation spectra obtained using a fast two-dimensional Laplace inversion. J Magn Reson 2002;154(2):261–268.

35. Venkataramanan L, Yi-Qiao S, Hurlimann MD. Solving Fredholm integrals of the first kind with tensor product structure in 2 and 2.5 dimensions. IEEE Transactions on Signal Processing 2002;50(5):1017–1026.

36. Macenko M, Niethammer M, Marron JS, Borland D, Woosley JT, Xiaojun G, Schmitt C, Thomas NE. A method for normalizing histology slides for quantitative analysis. 2009 28 June-1 July 2009. p 1107–1110.

37. Prasloski T, Rauscher A, MacKay AL, Hodgson M, Vavasour IM, Laule C, Madler B. Rapid whole cerebrum myelin water imaging using a 3D GRASE sequence. Neuroimage 2012;63(1):533–539.

38. Oshio K, Feinberg DA. GRASE (Gradient- and spin-echo) imaging: a novel fast MRI technique. Magn Reson Med 1991;20(2):344–349.

39. Does MD, Gore JC. Rapid acquisition transverse relaxometric imaging. J Magn Reson 2000;147(1):116–120.

40. Prasloski T, Madler B, Xiang QS, MacKay A, Jones C. Applications of stimulated echo correction to multicomponent T2 analysis. Magn Reson Med 2012;67(6):1803–1814.

41. Bonny JM, Traore A, Bouhrara M, Spencer RG, Pages G. Parsimonious discretization for characterizing multi-exponential decay in magnetic resonance. NMR Biomed 2020;33(12):e4366.

42. Doucette J, Kames C, Rauscher A. DECAES - DEcomposition and Component Analysis of Exponential Signals. Z Med Phys 2020;30(4):271–278.

43. Zlotzover S, Omer N, Radunsky D, Stern N, Blumenfeld-Katzir T, Reichman DB, Shrot S, Hoffmann C, Ben-Eliezer N. Improved myelin water imaging using B (1) (+) correction and data-driven global feature extraction: Application on people with MS. Imaging Neurosci (Camb) 2024;2.

44. Laule C, Kozlowski P, Leung E, Li DK, Mackay AL, Moore GR. Myelin water imaging of multiple sclerosis at 7 T: correlations with histopathology. Neuroimage 2008;40(4):1575–1580.

45. Benjamini D, Priemer DS, Perl DP, Brody DL, Basser PJ. Mapping astrogliosis in the individual human brain using multidimensional MRI. Brain 2023;146(3):1212–1226.

46. Endt S, Engel M, Naldi E, Assereto R, Molendowska M, Mueller L, Mayrink Verdun C, Pirkl CM, Palombo M, Jones DK, Menzel MI. In Vivo Myelin Water Quantification Using Diffusion-Relaxation Correlation MRI: A Comparison of 1D and 2D Methods. Appl Magn Reson 2023;54(11-12):1571–1588.

47. Kundu S, Barsoum S, Ariza J, Nolan AL, Latimer CS, Keene CD, Basser PJ, Benjamini D. Mapping the individual human cortex using multidimensional MRI and unsupervised learning. Brain Commun 2023;5(6):fcad258.

48. Garnier-Crussard A, Cotton F, Krolak-Salmon P, Chetelat G. White matter hyperintensities in Alzheimer’s disease: Beyond vascular contribution. Alzheimers Dement 2023;19(8):3738–3748.

49. Wardlaw JM, Valdes Hernandez MC, Munoz-Maniega S. What are white matter hyperintensities made of? Relevance to vascular cognitive impairment. J Am Heart Assoc 2015;4(6):001140.

50. Marstrand JR, Garde E, Rostrup E, Ring P, Rosenbaum S, Mortensen EL, Larsson HB. Cerebral perfusion and cerebrovascular reactivity are reduced in white matter hyperintensities. Stroke 2002;33(4):972–976.

51. Silbert LC, Roese NE, Krajbich V, Hurworth J, Lahna D, Schwartz DL, Dodge HH, Woltjer RL. White matter hyperintensities and the surrounding normal appearing white matter are associated with water channel disruption in the oldest old. Alzheimers Dement 2024;20(6):3839–3851.

52. Bettcher BM, Olson KE, Carlson NE, McConnell BV, Boyd T, Adame V, Solano DA, Anton P, Markham N, Thaker AA, Jensen AM, Dallmann EN, Potter H, Coughlan C. Astrogliosis and episodic memory in late life: higher GFAP is related to worse memory and white matter microstructure in healthy aging and Alzheimer’s disease. Neurobiol Aging 2021;103:68–77.

53. McAleese KE, Miah M, Graham S, Hadfield GM, Walker L, Johnson M, Colloby SJ, Thomas AJ, DeCarli C, Koss D, Attems J. Frontal white matter lesions in Alzheimer’s disease are associated with both small vessel disease and AD-associated cortical pathology. Acta Neuropathol 2021;142(6):937–950.

54. Eldeniz C, Finsterbusch J, Lin W, An H. TOWERS: T-One with Enhanced Robustness and Speed. Magn Reson Med 2016;76(1):118–126.

55. Leppert IR, Andrews DA, Campbell JSW, Park DJ, Pike GB, Polimeni JR, Tardif CL. Efficient whole-brain tract-specific T(1) mapping at 3T with slice-shuffled inversion-recovery diffusion-weighted imaging. Magn Reson Med 2021;86(2):738–753.

56. Dvorak AV, Kumar D, Zhang J, Gilbert G, Balaji S, Wiley N, Laule C, Moore GRW, MacKay AL, Kolind SH. The CALIPR framework for highly accelerated myelin water imaging with improved precision and sensitivity. Sci Adv 2023;9(44):eadh9853.

57. Dvorak AV, Wiggermann V, Gilbert G, Vavasour IM, MacMillan EL, Barlow L, Wiley N, Kozlowski P, MacKay AL, Rauscher A, Kolind SH. Multi-spin echo T(2) relaxation imaging with compressed sensing (METRICS) for rapid myelin water imaging. Magn Reson Med 2020;84(3):1264–1279.

58. Shin HG, Oh SH, Fukunaga M, Nam Y, Lee D, Jung W, Jo M, Ji S, Choi JY, Lee J. Advances in gradient echo myelin water imaging at 3T and 7T. Neuroimage 2019;188:835–844.

59. Xu G, Zhao Z, Zhu Q, Zhu K, Zhang J, Wu D. Myelin water imaging of in vivo and ex vivo human brains using multi-echo gradient echo at 3 T and 7 T. Magn Reson Med 2025;93(2):803–813.

60. Raman MR, Shu Y, Lesnick TG, Jack CR, Kantarci K. Regional T(1) relaxation time constants in Ex vivo human brain: Longitudinal effects of formalin exposure. Magn Reson Med 2017;77(2):774–778.

61. Shatil AS, Uddin MN, Matsuda KM, Figley CR. Quantitative Ex Vivo MRI Changes due to Progressive Formalin Fixation in Whole Human Brain Specimens: Longitudinal Characterization of Diffusion, Relaxometry, and Myelin Water Fraction Measurements at 3T. Front Med (Lausanne) 2018;5:31.

62. Seifert AC, Umphlett M, Hefti M, Fowkes M, Xu J. Formalin tissue fixation biases myelin-sensitive MRI. Magn Reson Med 2019;82(4):1504–1517.

63. Fritz FJ, Streubel T, Mordhorst L, Lüthi N, Edwards LJ, Mushumba H, Püschel K, Weiskopf N, Kirilina E, Mohammadi S. Longitudinal whole-human-brain quantitative MRI study on autolysis, fixation, rehydration, and shrinkage effects. bioRxiv 2026:2026.2001.2031.702882.

64. Shepherd TM, Flint JJ, Thelwall PE, Stanisz GJ, Mareci TH, Yachnis AT, Blackband SJ. Postmortem interval alters the water relaxation and diffusion properties of rat nervous tissue--implications for MRI studies of human autopsy samples. Neuroimage 2009;44(3):820–826.

65. Birkl C, Langkammer C, Golob-Schwarzl N, Leoni M, Haybaeck J, Goessler W, Fazekas F, Ropele S. Effects of formalin fixation and temperature on MR relaxation times in the human brain. NMR Biomed 2016;29(4):458–465.

66. Wang L, Oishi N, Urayama SI, Hanakawa T. Effects of Postmortem Intervals on Quantitative MRI in Unfixed and Fixed Swine Brain: Implications for Ex Vivo MRI Applications. Magn Reson Med Sci 2026.

67. Kim KW, MacFall JR, Payne ME. Classification of white matter lesions on magnetic resonance imaging in elderly persons. Biol Psychiatry 2008;64(4):273–280.

